# Induced mutagenesis by the DNA cytosine deaminase APOBEC3H Haplotype I protects against lung cancer

**DOI:** 10.1101/2020.02.28.970509

**Authors:** Mark A. Hix, Lai Wong, Ben Flath, Linda Chelico, G. Andrés Cisneros

## Abstract

The DNA cytosine deaminases APOBEC3A, APOBEC3B, and APOBEC3H Haplotype I can induce mutations in cells that lead to cancer evolution. The database of cancer biomarkers in DNA repair genes (DNArCdb) identified a single nucleotide polymorphism (rs139298) of APOBEC3H Haplotype I that is involved in lung cancer^1^. Here, we show that this single nucleotide polymorphism causes the destabilization of APOBEC3H Haplotype I. Computational analysis suggests that the resulting K121E change affects the structure of APOBEC3H leading to active site disruption and destabilization of the RNA-mediated dimer interface. A K117E mutation in a K121E background stabilized the APOBEC3H Haplotype I, enabled biochemical study, and showed that the K121E affected catalytic activity, single-stranded DNA binding, and oligomerization on single-stranded DNA. That the destabilization of a DNA mutator would be associated with lung cancer suggests that too much mutation could result in immune recognition or death of tumor cells, suggesting that multiple APOBEC3s would not be expressed in the same tumor cells. In support of this hypothesis, stably expressed APOBEC3H Haplotype I caused a high amount of double-stranded DNA breaks in a lung cancer cell line that endogenously expressed APOBEC3B. Altogether, the data support the model that high APOBEC3 mutations in tumors are protective.

## Introduction

The eleven member Apolipoprotein B editing enzyme (APOBEC) family in humans are comprised of RNA and single-stranded (ss) DNA cytosine deaminases with diverse biological functions^2-4^. Within this family are a subgroup of seven APOBEC3 family members that primarily deaminate cytosine in ssDNA, which forms promutagenic uracil. This C-to-U conversion is considered a DNA lesion and can result in a C-to-T mutation if uracil is used as a template during DNA replication or error prone uracil repair can nucleate other mutations, such as C-to-G transversions^5^. These enzymes are known for their ability to restrict the replication of various viruses, such as HIV-1, Epstein Barr Virus, and Hepatitis B virus, through the mutagenic fates of uracil in DNA^6-8^. The host genomic DNA is usually safe from cytosine deamination due to various protections such as cytoplasmic localization, low expression levels, or cytoplasmic RNA binding which both sequesters and inhibits enzyme activity^9-12^. However, when these checks are not in place, the APOBEC3 enzymes that can enter the nucleus, such as APOBEC3B (A3B), APOBEC3A (A3A), and APOBEC3H haplotype I (A3H Hap I)^9,13-18^, can deaminate genomic DNA. This activity has linked these APOBEC3 enzymes to various cancers, such as lung, breast, bladder, cervical, and others. The activity related to cancer is on transient genomic ssDNA during replication or transcription and is considered “off-target” activity^19-23^. Notably, the C-to-U conversion only occurs in a specific sequence context, e.g., 5’ RTCA (A3B, “R” is A or G), 5’ YTCA (A3A, “Y” is C or T), 5’TCT (A3H)^13,17,18^. This 5’TC sequence context has enabled the APOBEC3 mutation signature in at least 16 cancers and can be differentiated from chemical DNA damage, which may occur with lung cancer, for example^5,24^. Despite significant characterization of APOBEC3-induced mutations in tumors, it is not known if APOBEC3-induced mutations can initiate a cancer, cause evolution of a cancer or both^25^. The main hypothesis is that APOBEC3 enzymes create uracils randomly throughout the genome and that the majority of these are repaired by Base Excision Repair, but some may not be repaired, resulting in transition mutations, and some may be repaired in an error prone manner, resulting in transversion mutations^26-28^. Too few mutations and there is no effect, too many mutations and there is cell death or immune recognition, and somewhere in the middle, APOBEC3 enzymes can cause a “just right” rate of mutations for tumor evolution^28^. These fates depend on the cell cycle and the numbers of mutations. Although this mutagenic process can make up at least 60% of the mutations in a tumor genome, the mutagenesis is random and acts to increase the genetic diversity in the tumor cell population, facilitating the chance of a beneficial mutation arising that helps maintain the cell population^5,13,17,27,29,30^. The interplay between APOBEC3 enzymes in this process is not known. Although there have been multiple reports of A3B, A3A, or A3H Hap I individually being involved in a cancer, it is not known if more than one APOBEC3 is expressed in a cancer cell at the same time and what would be the effects^13,16-18,31^. What has recently been identified is that APOBEC3 mutations occur episodically in cancers^32^. This is thought to be because constant expression and mutagenesis would result in cell death or recognition by the immune system, rather than creating a “just right” level of mutagenesis for cancer evolution. It is also known that for lung cancer, A3H Hap I predominantly causes early mutations and A3B predominantly causes late stage mutations^13^.

Interestingly, there are seven haplotypes for A3H, defined by polymorphisms or deletions at positions 15, 18, 105, 121, and 178, but only A3H Hap I is primarily localized to the nucleus^33-35^. The other A3H haplotypes are primarily in the cytoplasm^35^. Only those APOBEC3 members that have access to the nucleus can contribute to cancer mutagenesis. In addition, not all A3H haplotypes are catalytically active. Some haplotypes are unstable (III, IV, VI) some are hypo-stable (I) and others are stable (II, V, VII)^33-35^. Despite A3H Hap I being hypo-stable/-active, it is the only A3H that has been implicated in cancer mutagenesis. Starrett *et al.* have shown that A3H Hap I is an enzymatic contributor to ‘APOBEC signature’ mutations in lung cancer and likely contributor to breast cancer^13^. This was a surprising result since A3H Hap I has a half-life in cells of approximately 30 min^33^. The short A3H Hap I half-life also precludes the purification of significant quantities of A3H Hap I for use in biochemical assays. When cell lysates are used and A3H Hap I is overexpressed, at equivalent protein levels to more stable haplotypes, the activity is comparable^13^. A3H Hap I has a G105 that causes the low stability and mutation of this to an R105 forms a stable enzyme and is A3H Hap VII, which has been used as an A3H Hap I proxy *in vitro*^*36*^.

One single nucleotide polymorphism, rs139298, resulting in A3H Hap I K121E has been recently reported to be associated with lung cancer^1^. Lung cancer results in over 1.6 million deaths per year and nearly 2 million new cases annually, with pulmonary adenocarcinoma comprising about 40% of the total cases of non-smoking lung cancer^37-40^. Surprisingly, we found that this K121E mutation further destabilized A3H Hap I, giving the counter-intuitive result that the absence of A3H Hap I is associated with lung cancer. The data in the present work support the hypothesis that although APOBEC3 enzymes contribute to cancer mutagenesis, constant exposure to APOBEC3 mutagenesis or multiple APOBEC3s may be detrimental. In this study, we examine the A3H Hap I K121E variant and its stabilization by a novel A3H Hap I stabilizing mutation, K117E, using computational, biochemical, and cellular techniques. The combined results demonstrate that the timing of APOBEC3 mutations and the number of different APOBEC3s jointly expressed in cancer cells influences the fate of APOBEC3-induced somatic mutagenesis and provides the first cause and effect evidence of APOBEC3 enzymes and their role in cancer mutagenesis.

## Results

### A3H Hap I K121E results in the formation of a new hydrogen-bonding network

We used classical molecular dynamics (MD) to investigate the effect (if any) of the SNP resulting in the A3H Hap I K121E mutation on the protein structure and dynamics compared with A3H Hap I wild type (WT). Pairwise hydrogen bond analysis of the K121E indicates the formation of a new hydrogen bond (HBond) network across part of the protein surface compared with the WT. This HBond network induces strain on the active site suggesting a change in stability or activity (**Figure 1**). Hydrogen bond interactions between protein residues that differed by more than 30% of the simulation time were identified. The HBond network in the K121E system connects two existing networks (R124 to I182 and K117 to P118 to S87) from the WT into a single larger network in the K121E variant. This extended over-stabilized network effectively “freezes” the helix where the mutation occurs, disrupting both the active site and RNA-binding residues on the C-terminal helix. Changes in root mean squared fluctuation (RMSF) are also observed in the K121E system, with residues involved in the HBond network exhibiting reduced fluctuation while the rest of the protein exhibits increased fluctuation (**Figure 2**). Principle component analysis (PCA) shows that the first two modes of motion in the K121E system are constrained and tightly correlated in comparison to the WT (**Figure 2**). Difference correlation matrices reveal that K121E has large regions of changing correlated movement, especially with respect to the region around the mutation point.

**Figure 1:**
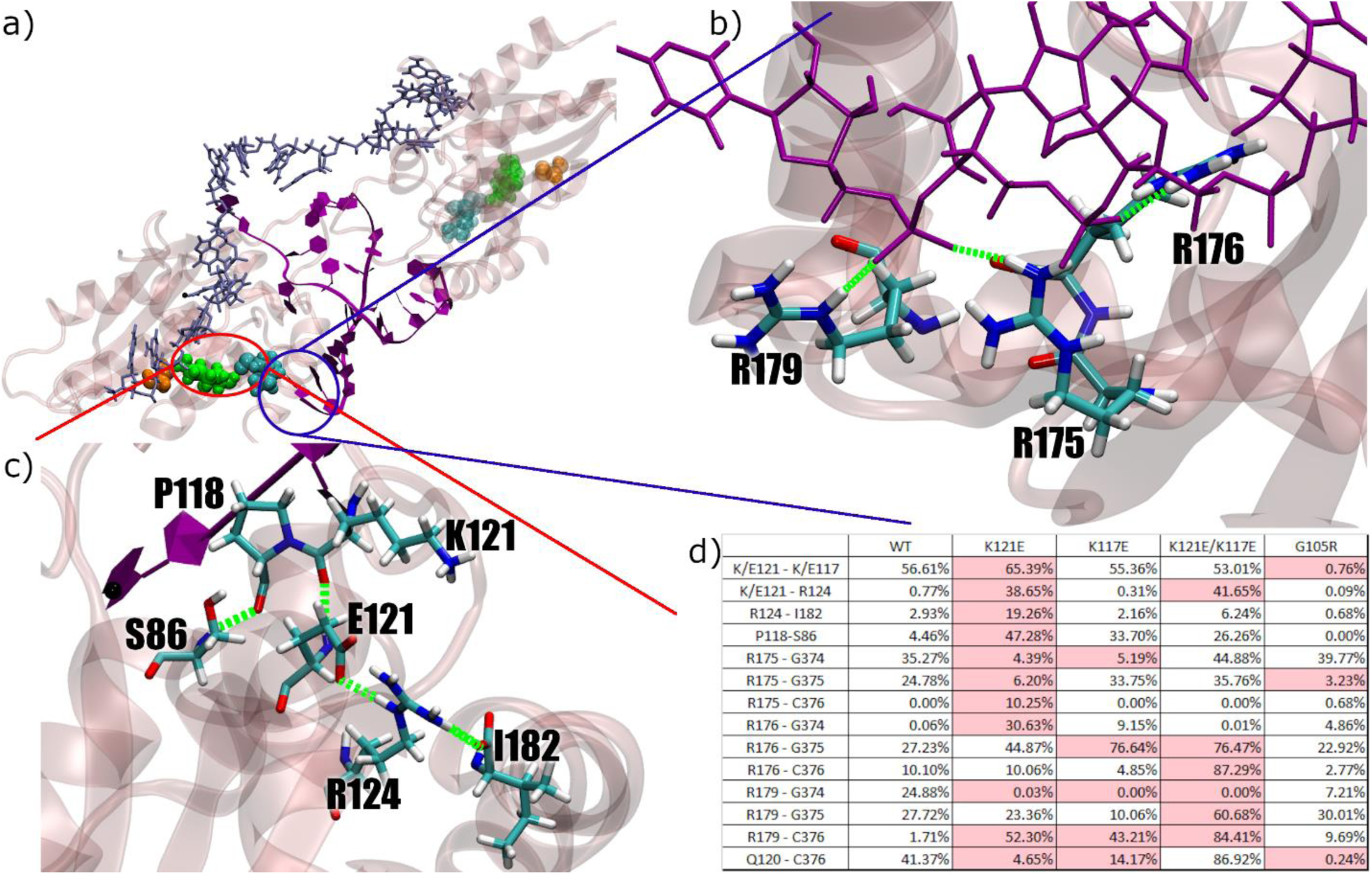
**a)** APOBEC3H protein (transparent pink) with RNA interface (purple) and DNA substrate (iceblue). G105 is shown in orange, K117 in cyan, and K121 in green, with corresponding residues on opposite monomer shown in transparency. **b)** RNA-binding residues on terminal helix. **c)** Formed hydrogen bonding network as result of K121E mutation. **d)** Table of largest contributors in each hydrogen bonding interaction, shown as percentage of total simulation time.

**Figure 2:**
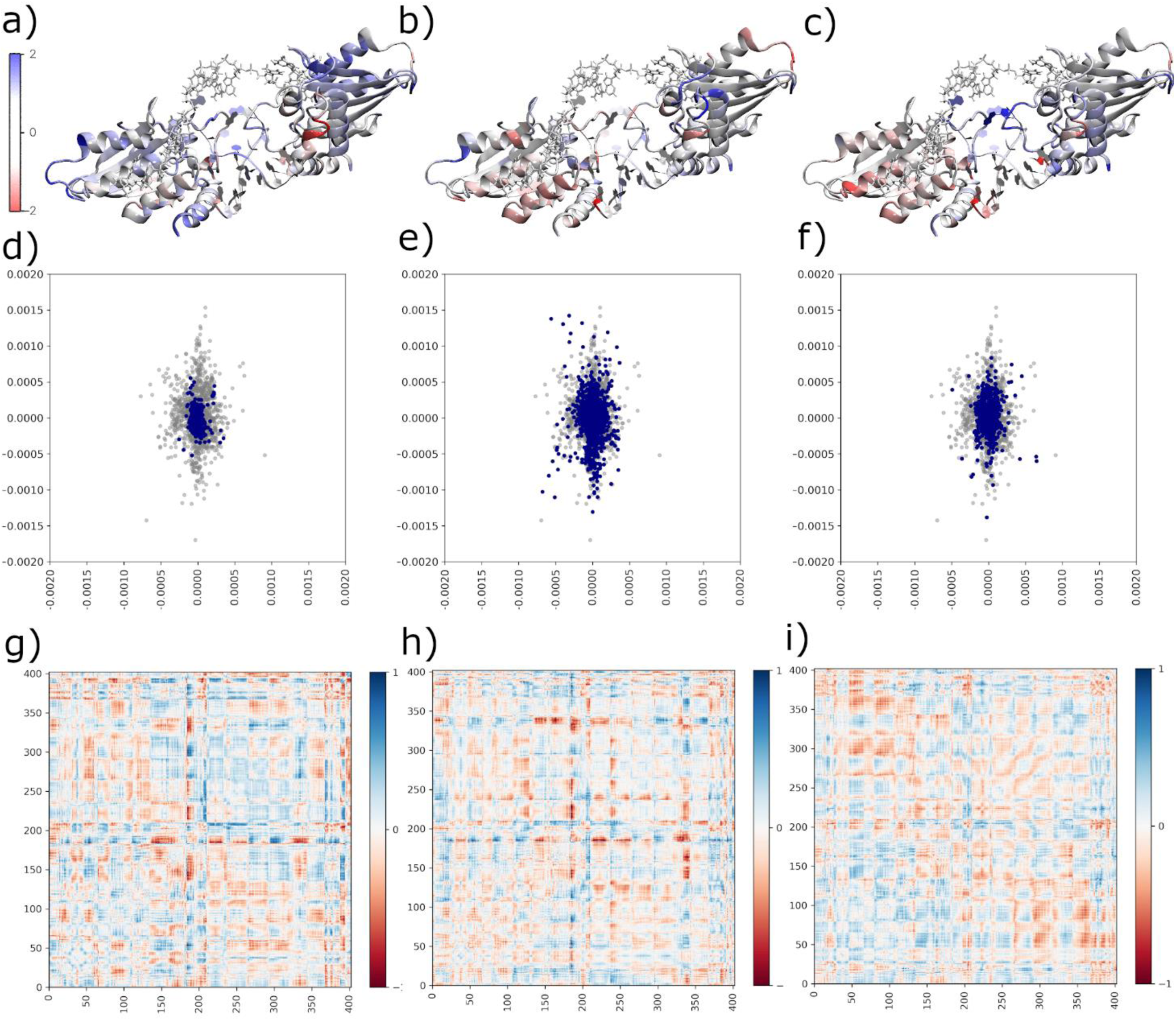
Difference RMSF with respect to wildtype overlaid on protein structures for **a)** K121E, **b)** K121E/K117E, and **c)** K117E. Principle component analyses for mutant systems (blue) over wildtype (grey) for **d)** K121E, **e)** K121E/K117E, and **f)** K117E. Difference correlation matrices against wildtype reference for **g)** K121E, **h)** K121E/K117E, and **i)** K117E.

The simulation data suggest that a K117E mutation could rescue the K121E variant by restoring the two hydrogen bonding networks. The K121E/K117E variant removes the HBond between the side chains of 121 and 117, and partially restores the remaining interactions in the network towards their WT levels. (**Figure 1**). The K121E/K117E system shows less of a change in fluctuation, with residues in the network exhibiting near WT fluctuation, similarly to what is observed for G105R (**Figure 2**). The PCA shows that the first two modes are similar to those of the WT, as does the comparatively minor change in correlated movement.

### K121E destabilizes A3H Hap I

To test the computational results we produced the A3H Hap I variants in 293T cells for cell-based analysis. To determine the steady state expression levels of the A3H Hap I WT, K121E cancer variant, putative rescue mutant K117E/K121E, and rescue mutant control K117E, we transfected 3x HA-tagged A3H expression constructs into 293T cells. Immunoblotting showed that steady state expression levels for the cancer variant were not detectable by immunoblotting, indicating that the K121E mutation destabilized A3H Hap I WT (**Figure 3a**). We found that the K117E mutation stabilized A3H Hap I in the presence of the otherwise destabilizing G105 amino acid^33^ and, as predicted, can stabilize the cancer variant (K117E/K121E) (**Figure 3a**). Since the destabilizing SNP was associated with lung cancer^1^, these data suggested that A3H Hap I somatic mutations may be detrimental to cancer progression perhaps due to extensive mutagenesis leading to cell dysfunction or immune recognition. This is consistent with A3H Hap I mutations being identified early in cancer and the episodic nature of APOBEC3-induced mutations in cancer^1,13,32^.

**Figure 3:**
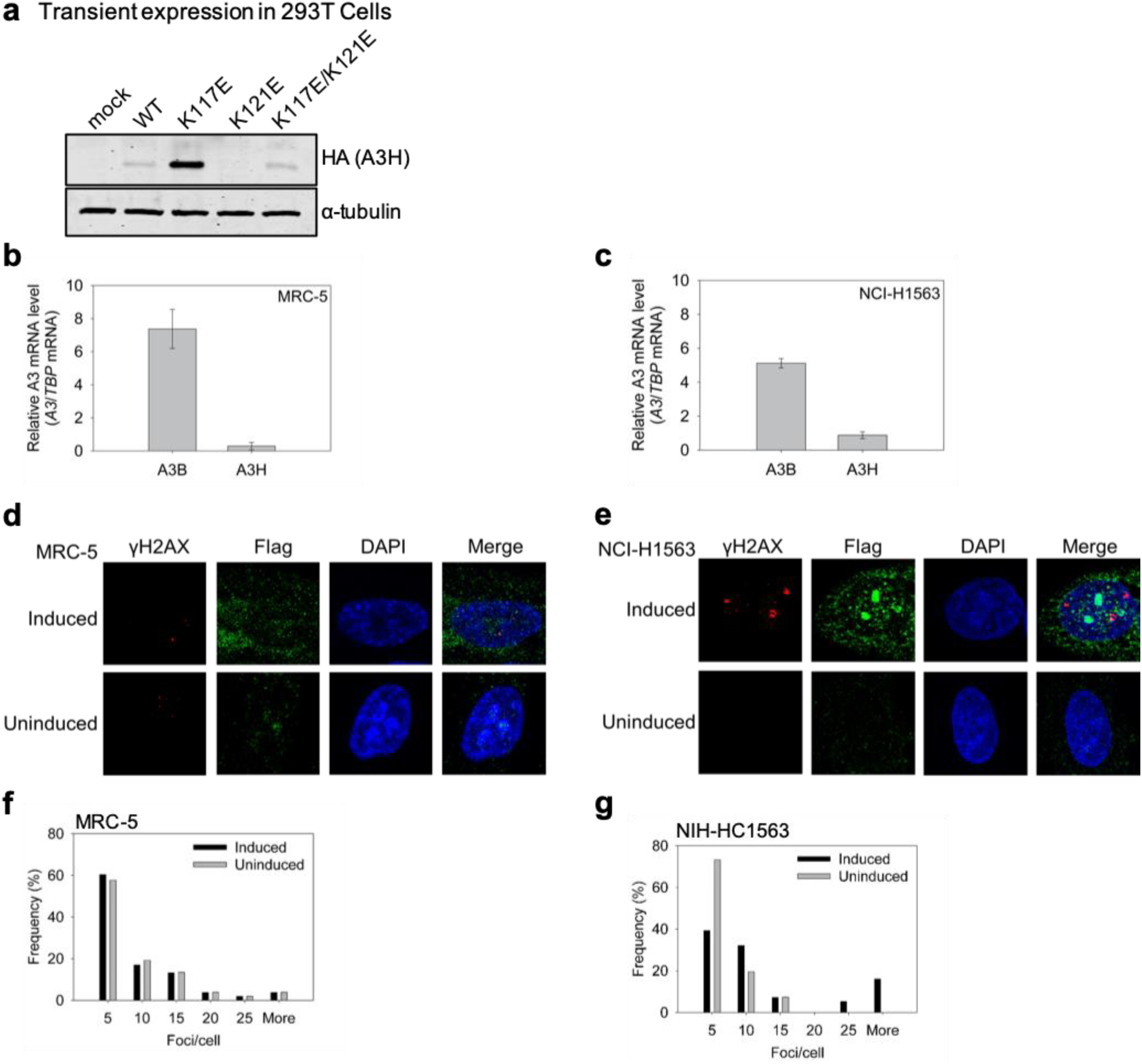
Expression and mutagenic activity of A3H Hap I. **a)** Transient expression of HA-tagged A3H Hap I expression constructs in 293T cells were detected by immunoblotting. The α-tubulin served as the loading control. The endogenous *A3B* and *A3H Hap I* mRNA levels were detected in **b)** MRC-5 and **c)** NCI-H1563 cells relative to *TBP* mRNA. The A3H Hap I mRNA levels were low, so doxycycline inducible A3H Hap I-Flag cell lines were produced (Figure SF6). **(d-g)** A3H Hap I induces high amounts of γH2AX foci in NCI-H1563 cells with endogenous A3B expression. The γH2AX foci were detected in induced and uninduced cells by immunofluorescence microscopy in **d)** MRC-5 and **e)** NCI-H1563 cells. Results were quantified and plotted as a histogram for **f)** MRC-5 and **g)** NCI-H1563 cells.

### Combined expression of A3B and A3H Hap I in a lung cancer cell line induces high amounts of double-stranded DNA breaks

To directly test this hypothesis, we used lung cell lines from two sources representing normal (MRC-5, male fetal tissue) and cancer (NCI-H1563, adenocarcinoma from a male non-smoker) cells. We first determined if these cell lines expressed any endogenous A3s. We found that both the MRC-5 and NCI-H1563 cell lines expressed high amounts of A3B mRNA compared to the TBP mRNA control (**Figure 3b-c**). The two cell lines expressed 7-fold (MRC-5) and 5-fold (NCI-H1563) more *A3B* mRNA than *TBP* mRNA, which is equivalent to the amounts of *A3B* mRNA found in breast cancer cell lines and responsible for genomic mutations^17^. The endogenous *A3H* (not genotyped) was at an equivalent level to *TBP* and 7- to 5-fold less than A3B (**Figure 3b-c**). This provided an opportunity to transduce these cells with an A3H-Hap I-Flag expression construct to make doxycycline (dox)-inducible stable cell lines (**Figure SF6**) and test the effect of multiple A3s expressed in normal and cancer lung cells. A key marker of high A3 activity is the formation of double-stranded (ds) DNA breaks that are induced by base excision of closely located uracils. To avoid any possible cell death, we only induced cells with dox for 24 h, then cells were fixed and antibodies used to detect γH2AX foci as a marker of dsDNA breaks.

In MRC-5 cells the endogenous A3B alone and A3B with A3H-Hap I-Flag were not significantly different and showed that ∼60% of cells had five γH2AX foci/cell, ∼20% of cells had ten γH2AX foci/cell, and ∼13% of cells had fifteen γH2AX foci/cell. Larger numbers of foci per cell were observed in less than 10% of the population (**Figure 3d, f**). These data support that there is sufficient DNA repair to process A3-induced dsDNA breaks from multiple A3s in normal cells.

In NCI-H1563, the lung adenocarcinoma cell line, there are mutations in KRAS, PIK3R1, SMARCA4, and p53^41^, indicating that it has common cancer mutations that dysregulate cell cycle, cell division and chromatin remodelling. This makes cancer cells more susceptible to A3-induced mutagenesis. However, with only endogenous A3B expressed, the number of γH2AX foci/cell were approximately the same as both conditions tested for MRC-5 cells, i.e., ∼60 to 70% of cells having five γH2AX foci/cell (**Figure 3e, g**). Nonetheless, when A3H Hap I-Flag expression was induced the number of γH2AX foci/cell increased dramatically with 16% of cells having over twenty-five γH2AX foci/cell (range was between 33 to 61 per cell). The other major populations were 32% and 39% of cells with ten and five γH2AX foci/cell, respectively (**Figure 3e, g**). These data demonstrate that more than one A3 expressed in a cancer cell can induce a large amount of dsDNA breaks, which may result in cell death or chromosomal aberrations. This supports the genetic data that the loss of A3 activity, specifically A3H Hap I, can be associated with cancer^1^.

### A G105R mutation results in a more active and stable A3H than a K117E mutation

Although a K117E mutation can stabilize A3H Hap I, it is not a naturally occurring variant. The naturally occurring stabilization of A3H Hap I is a G105R mutation, which is known as A3H Hap VII. The reason for the instability induced by G105 is not known. Computational models of the G105R system exhibit dynamic motion and hydrogen bonding patterns similar to the WT, indicating that this mutation does not negatively affect the structural stability of the protein (see SI for G105R computational data). The G105R may simply increase the propensity of the protein to become polyubiquitinated and degraded^42^, in contrast to K121E that destabilizes the structural integrity of A3H Hap I (**Figure 2**). To test if the longer steady state half-life of A3H K117E and K117E/K121E in cells correlates with increased catalytic activity similar to A3H Hap VII, we conducted an *in vitro* deamination assay using protein purified from *Sf*9 insect cells to obtain a quantitative measurement. A3H deaminase activity was observed by adding a 118 nt ssDNA with two A3H deamination motifs (5’CTC) and an internal fluorescein label to equivalent amounts of the cell lysates. The two 5’CTC motifs account for the preference of some A3H variants to preferentially deaminate one cytosine motif more than the other^43^. We found that despite recovery of protein stability the A3H K117E and K117E/K121E were 4-fold less active than A3H Hap VII (relative to Hap I, G105R) or A3H Hap II (relative to Hap I, G105R, E178D) (**Figure 4a**). These data suggest that the WT hydrogen bonding network is important for catalytic activity and compensatory mutations can correct the instability, but not fully reinstate catalytic activity. Despite the decreased specific activity, the mutants were also processive, similar to what has been reported for A3H Hap VII and A3H Hap II (**Figure 4b**)^43^.

**Figure 4.**
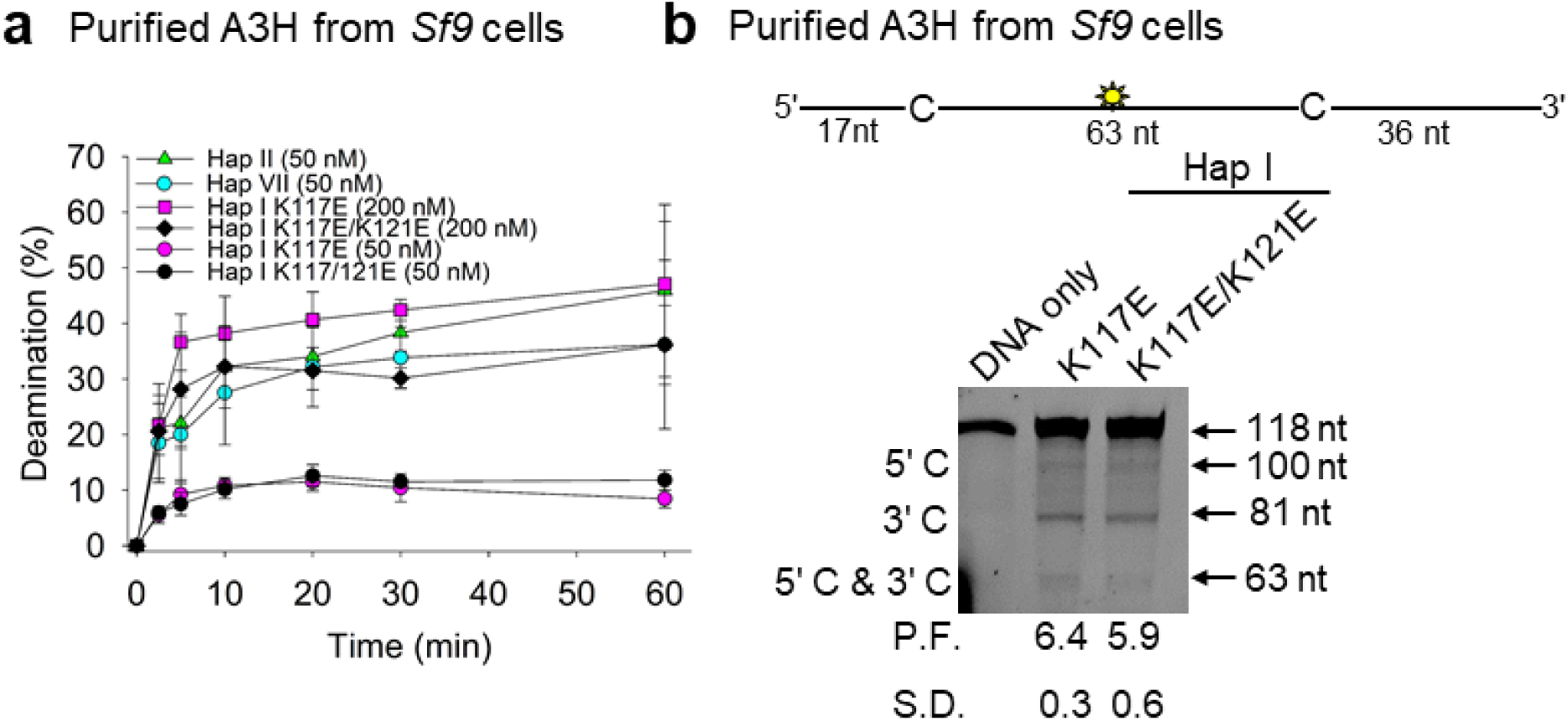
Deamination activity of A3H WT and mutants. **a)** Time course of deamination of A3H Hap II, A3H Hap VII, and A3H Hap I WT and mutants. Deamination was tested on a 118 nt ssDNA (100 nM) and gels analyzed for each plot are shown in Figure SF5. **b)** Processivity of A3H Hap I K117E and K117E/K121E. The processivity factor (P.F.) measures the likelihood of a processive deamination over a nonprocessive deamination and is described further in the Methods. Both K117E and K117E/K121E are approximately 6-fold more likely to undergo processive deamination than a nonprocessive deamination. The standard deviation for three independent experiments is shown as (**a**) error bars or (**b**) below the gel.

### A3H cancer variant destabilizes dimer interface

From monkeys to greater apes, the A3 family has lost activity. Although A3B activity can be lost through a gene deletion^44^, other A3s have lost activity because of decreased catalytic activity due to mutation of residues near the active site, e.g., A3D or loss of dimerization, e.g., A3C^45,46^. Based on recent crystal structures, the association of A3H with RNA is concomitant with enzyme stability and dimerization^9,47-50^. This is a unique feature of A3H compared to other A3s, in that it uses a double-stranded RNA molecule to form a dimer interface without any protein-protein contacts and that this imparts stability to the enzyme. As a result, we hypothesized that the reason for the A3H cancer variant instability (**Figure 3a**) may be due to the inability to bind RNA and dimerize. RNA-mediated dimerization can be detected by treating a purified protein preparation with RNase A and observing if a ∼12 nt RNA is protected from digestion. We confirmed that recombinant A3H Hap I mutants K117E and K117E/K121E purified from *Sf*9 cells with RNA and RNase A treatment resulted in a ∼12 nt RNA being protected from degradation by the protein (**Figure 5a**). The A3H Hap VII and A3H Hap II also bound cellular RNA and protected a ∼12 nt RNA in the presence of RNase A. Accordingly, all four A3H had the same Size Exclusion Chromatography (SEC) profile and formed primarily a dimer (**Figure 5b-d**).

**Figure 5:**
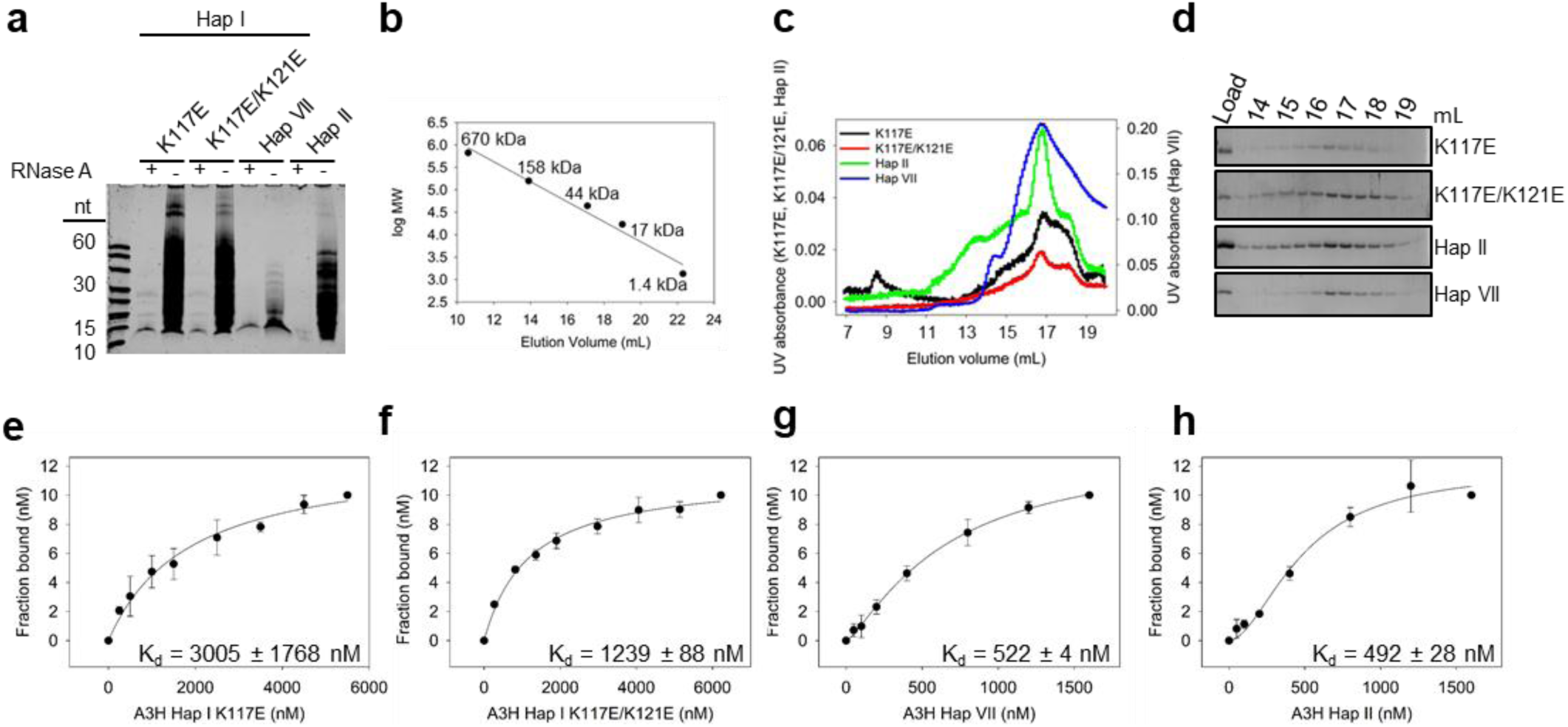
A3H Hap I mutant dimerization, oligomerization, and ssDNA binding. **a)** A3H dimerization is mediated by RNA. **a)** Purified A3H was denatured in formamide buffer and samples were resolved by urea denaturing PAGE. The gel was stained with SYBR Gold to detect nucleic acids. The A3H Hap I K117E, A3H Hap I K117E/K121E, A3H Hap VII, and A3H Hap II protect an approximately 12-15 nt RNA. Representative image shown from three independent experiments. **b)** Standard curve and **c-d)** SEC profile for A3H Hap I K117E, A3H Hap I K117E/K121E, A3H Hap VII, and A3H Hap II demonstrating similar elution profiles that are composed primarily of a dimer peak (44 kDa, 17 mL elution volume). **(e-h)** The apparent K_d_ of A3H enzymes from a 118 nt ssDNA was analyzed by steady-state rotational anisotropy. The **e)** A3H Hap I K117E (3005 ± 1768 nM) and **f)** A3H Hap I K117E/K121E (1239 ± 88 nM) bind the ssDNA with different affinities than **g)** A3H Hap VII (522 ± 4 nM) and **h)** A3H Hap II (492 ± 28 nM). The apparent K_d_ was calculated by determining the best fit by least squares analysis which was a hyperbolic fit for **e-f** and a sigmoidal fit for **g-h**. Error bars represent the S.D. from three independent experiments.

Since we cannot purify the K121E variant, we used computational analysis to predict whether the change at amino acid 121 would destabilize dimerization. RMSF analysis for WT, K121E and K117E/K121E (**Figure 2**) shows that residues in the K121E system exhibited increased fluctuation compared with WT, except those residues involved in the new hydrogen bonding network and many that interact with the RNA interface. These RNA-binding residues (R175, R176, R179) exhibit changing hydrogen bonding interactions with the RNA interface. In the simulations, these changes cause the RNA to change positions, affecting additional protein-RNA interaction at loop 1 residues. Correlation matrices were calculated for all systems, with the K121E variant system showing increased correlation between the monomer subunits and decreased correlation within each monomer when compared to the WT (**Figure SF2**), suggesting that the dimer interface is destabilized by the newly formed hydrogen bonding network. The K117E system shows qualitatively similar zones of correlation and anticorrelation at the dimer interface with respect to the WT. The K117E/K121E mutant correlation matrices reveal that this system more closely resembles the K121E than either the K117E or WT dimer systems, with qualitatively similar correlation matrices. These data are consistent with poor expression in cells (**Figure 3a**). Further, consistent with the biochemical data, the K117E/K121E variant system exhibits fewer changes in fluctuation, and there is not a general trend of increased fluctuation as seen in the K121E variant.

### A3H variants have weakened interactions with the substrate

We observed that both the A3H Hap I K117E and K117E/K121E had ∼6-fold (K117E) and ∼2.5-fold (K117E/K121E) higher apparent dissociation constants from ssDNA than A3H Hap VII or A3H Hap II (**Figure 5e-h**). Further, both the A3H Hap I K117E and K117E/K121E binding curves best fit to a rectangular hyperbola by least squares analysis, in contrast to A3H Hap VII and A3H Hap II that best fit to a sigmoid. The sigmoidal binding relationship for A3H Hap II has been shown to be due to further oligomerization of A3H dimers on ssDNA^48^. This does not appear to occur with A3H Hap I K117E or K117E/K121E (**Figure 5e-f**), although they can form dimers in solution (**Figure 5b-d**). The hindered oligomerization on ssDNA that may be due to a faster off-rate from the substrate likely is what affected the deamination activity (**Figure 4a**). Altogether, the data indicate that the SNP that inactivates A3H Hap I results in an inability to use RNA for dimerization, emphasizing the importance of cellular RNA to the structural stability of A3H. Further, the hydrogen bonding network is important for interactions with ssDNA and oligomerization on ssDNA, providing a reasoning for the decreased catalytic activity in A3H Hap I K117E and K117E/K121E.

## Discussion

A3H Hap I was identified to induce genomic mutations specifically in lung cancer^13^. Interestingly, the activity of A3H Hap I was found because researchers wanted to investigate how genomic mutations with a cytidine deaminase signature were still occurring in people with an *A3B*_*-/-*_ deletion^51-54^, which occurs in the world population at 22.5%, although it primarily occurs in Oceanic populations^44^. Although A3A is also involved in cancer mutagenesis, it appears to be strongly associated with breast cancer^31,55^. The association of A3H Hap I expression and mutation signature were strong in early clonal variants of lung cancers. Further supporting a role of A3H Hap I in lung cancer was a computational study that identified the association of SNP rs139298 in A3H Hap I in lung cancer^1^. However, we demonstrate in this work that the SNP makes the A3H Hap I completely unstable, suggesting that loss of this enzyme specifically promoted cancer. While the effects arising from the combination of environmental, e.g., smoking, and A3 mutations remain unknown, we did assess if multiple A3 enzymes would cause too many mutations and be detrimental to cancer evolution. Recent reports support the seemingly contradictory idea that A3 enzymes, specifically APOBEC3B can both contribute to cancer and contribute to success of cancer immune therapy due to A3-mutation created neoepitopes^56^ or efficiency of platinum-based drugs due to contributions to DNA damage^57^.

In this report we came to an equivalent conclusion since we determined that the previously identified rs139298 SNP that creates a K121E mutation in A3H Hap I resulted in a loss of enzyme stability in cells. While based on previous evidence, it appears that A3H Hap I, in an A3B null background, can promote genome mutations that contribute to lung cancer^13^, it had not been investigated what would occur in the presence of another A3. Surprisingly, we found that two lung cell lines, one normal and the other cancerous, expressed similar and high amounts of endogenous A3B. The A3H Hap I in the cancer cell line, co-expressed with A3B increased the number of dsDNA breaks significantly compared to A3B alone. These dsDNA breaks could lead to cell death, destructive chromosomal relocations or high amounts of neoantigens. While in many A3/cancer studies it is difficult to determine the fate of A3 mutations, the genetic data combined with computational and biochemical study of the K121E mutation demonstrates that A3H Hap I has these detrimental effects on cell growth and proliferation in lung cancer.

The inactivation of A3 enzymes through SNPs is not unique to A3H, but A3H does have the highest number of inactivating SNPs in the A3 family. In particular, A3H has been lost multiple times in primate evolution, presumably due to nuclear localization and acquisition of genomic mutations^33^. While it is thought that A3H is kept active at a certain level in the population due to the antiviral effects of the enzyme, there are strong population stratifications in A3H active/hypo-active/not active forms that correlate with the historical levels of HIV infection, a pathogen that active A3Hs can restrict efficiently^58,59^. The antiviral properties of these enzymes are likely why they are maintained despite the negative effect of their off-target activity. The destabilizing SNPs for A3H other than the one studied here, are not thought to destabilize protein structure, but to promote A3H ubiquitination and proteosomal degradation^42^. Thus, the SNP studied in this work for A3H Hap I is unique since it appears to destabilize the enzyme by disrupting a hydrogen bonding network. While this disruption decreased catalytic activity, it did not cause protein unfolding. Rather, it decreased the interaction strength with a dsRNA molecule that A3H acquires from the cell and uses to dimerize^9,47-50^. This dimerization using an RNA intermediate is essential to A3H thermodynamic stability^47,48^. While the nature of dimerization for different A3s is unique, e.g., only A3H uses an RNA intermediate, a SNP that regulates activity through dimerization has also been found for A3C^46,60^. This type of functional inactivation enables rapid evolutionary toggling of activity for times when enzyme activity is needed again^61^.

Since the identification of distinct mutational signatures and high A3 expression in cancers^5,16,17^, a number of research groups have contributed to our understanding of how A3 enzymes access ssDNA, which A3 enzymes are involved, and specifically for A3B, the clinical effects of the induced mutations^18,19,21,23,62-64^. However, there are no studies examining multiple A3 expression in a cancer cell, although this potentially could happen with three A3 enzymes involved in cancer mutagenesis. The data presented here supports the idea that the expression of multiple A3s in a single cell may benefit the host and be detrimental to the cancer since the A3H Hap I K121E is associated with lung cancer, and as shown here, is destabilized in cells. Based on analysis of dsDNA breaks, it is likely that at least A3H Hap I and A3B are not compatible for promoting cancer due to the accumulation of too many mutations. In combination with the genetic data^1^, our data provide direct evidence for such a cause and effect relationship of A3s and cancer.

## Methods

### Computational Methods

Molecular dynamics (MD) simulations were performed with the ff14SB^65^, OL15^66^, YIL^67^, and TIP3P^68^ force fields using the pmemd.cuda^69,70^ and OpenMM^71^ programs. All systems were constructed from pdbid 5W45, corresponding to A3H-HapI based on the UniProt consensus sequence^72,73^. A search of 12 possible binding orientations of the DNA substrate on one single monomer was performed to determine the most probable orientation for the DNA substrate (see SI for coordinates of dimer in holo form)^74^.

The most probable orientation was subsequently used to construct the biologically relevant dimer systems using the macaque-APOBEC3H dimer structure (pdbid 5W3V) as a template for UCSF Chimera and taking into account the polarity of the substrate^75-77^. The monomer crystal of human A3H was overlaid on the dimer crystal of the macaque A3H and aligned. The protonation states for the dimer systems were determined using H++, neutralized with K^+^ (**Table ST2**) and solvated in a box of TIP3P water of approximately 143Å on each side to allow free movement without self-interaction across the periodic boundary^78-80^. Particle mesh Ewald was employed for long-range electrostatics, with a cutoff distance of 10 Å and an error tolerance of 10^−4^ kJ mol^-1^ Å^-2^.^70^

All simulations were performed in the NPT ensemble at 1.0 bar and 300 K using a Monte Carlo barostat, and a Langevin thermostat with a 1 fs timestep^71,81,82^. Simulation frames were saved at 10 ps intervals. To maintain active site geometry in the simulations, distance restraints of 20 kcal/mol· Å^2^ at 2.0 Å were applied between the Zn^2+^ atoms and their respective coordinating histidines, with additional 10 kcal/mol· Å^2^ restraints at 5.0 Å between the Zn^2+^ and water-coordinating glutamate to prevent incursion into the active site. Each system was minimized for 2,000 steps and equilibrated for 20ns before beginning production. Additional simulations using distance constraints of 2.0 Å between the Zn^2+^ and coordinating atoms were also run and found to give quantitatively similar results to the restrained simulations (data not shown).

Five systems were constructed including A3H Hap I wildtype (WT), A3H Hap I K121E (cancer variant), A3H Hap I K117E/K121E, A3H Hap I K117E, and A3H Hap I G105R. Every system was run for 250 ns in triplicate (750ns total simulation time per variant). Energy decomposition analysis (EDA) was performed using an in-house FORTRAN90 program^83-85^. RMSF, RMSD, correlation, hydrogen bonding analysis, clustering, normal mode analysis, and distance calculations were performed using cpptraj in the AmberTools suite^86,87^.

### Immunoblotting

Transfection of 1 × 10^5^ 293T cells per well of a 6-well plate was carried out using GeneJuice (Novagen) transfection reagent as per manufacturer’s protocol. Cells were maintained in DMEM with 10% FBS. Plasmids transfected were empty pcDNA3.1 (mock) and the following C-terminally tagged A3H-3x HA constructs in pcDNA3.1: A3H Hap I, A3H Hap I K117E, A3H Hap I K121E, and A3H Hap I K117E/K121E. After 48 h, cells were lysed using 2x Laemmli Buffer and 30 μg total protein was used. A3H was detected using an anti-HA mouse antibody (Sigma) and loading control for cell lysate detected α-tubulin using an anti-α-tubulin rabbit antibody (Invitrogen). Secondary detection was performed using Licor IRDye antibodies produced in goat (IRDye 680-labeled anti-rabbit and IRDye 800-labeled anti mouse).

### Purification of A3H from *Sf9* insect cells

The pFAST-bac1vectors were used to produce recombinant baculovirus according to the protocol for the Bac-to-Bac system (Life Technologies). Recombinant GST-A3H Hap I WT and mutant baculovirus (K117E, K117E/K121E) was used to infect *Sf9* cells at a multiplicity of infection of 1 and cells were harvested after 72 h. Cells lysates were treated with 100 μg/mL of RNase A (Qiagen). Lysates were cleared by centrifugation and then incubated with Glutathione-Sepharose 4B resin (GE Healthcare) at 4 °C overnight and subjected to a series of salt washes (0.25M to 1M NaCl) as described previously^88^. On-column cleavage from the GST tag with Thrombin (GE Healthcare) was performed at 21 °C for 18 h in thrombin digestion buffer (20mM HEPES, pH 7.5, 150 mM NaCl, 10% glycerol, and 2 mM DTT). Proteins were assessed to be 90% pure by SDS-PAGE (**Figure SF7**).

### Deamination assays

The 118 nt ssDNA substrate was previously described^43^. The 118 nt ssDNA substrate had the following sequence: 5′GAA TAT AGT TTT TAG CTC AAA GTA AGT GAA GAT AAT [Fluorescein-dT] TAG AGA GTT GTA ATG TGA TAT ATG TGT ATG AAA GAT ATA AGA CTC AAA GTG AAA AGT TGT TAA TGT GTG TAG ATA TGT TAA. The ssDNA substrate (100 nM), which contained two A3H deamination motifs (5′CT<u>C</u>, with the underlined C being deaminated), was incubated with 200 nM of A3H Hap I mutants (K117E, K117E/K121E) or 50 nM of A3H Hap II, Hap VII and Hap I mutants for 60 min at 37 °C in buffer that contained (50 mM Tris pH 7.5, 40 mM KCl, 10 mM MgCl_2_, and 1 mM DTT). For determination of enzyme processivity, additional reactions were carried out under single-hit conditions, i.e., <15% substrate usage, to ensure that deaminations on each ssDNA were catalyzed by a single enzyme^89^. Under these conditions, a processivity factor can be determined by comparing the total amount of deaminations occurring at two sites on same ssDNA to a calculated theoretical value of deaminations at these two sites if the deamination events were uncorrelated (not processive)^90^. A3H catalyzed deaminations were detected by treating the ssDNA with Uracil DNA Glycosylase (New England Biolabs) and heating under alkaline conditions before resolving the Fluorescein dT-labeled ssDNA on a 10% (v/v) denaturing polyacrylamide gel. Gel images were obtained using a Chemidoc-MP imaging system (Bio-Rad) and integrated gel band intensities were analyzed using ImageQuant (GE Healthcare).

### Detection of RNA bound to A3H

To examine the RNA present in purified A3H, 3.5 μg A3H was or was not treated with RNase A (Roche Applied Science) for 15 min at room temperature. At the end of the reactions, samples were mixed with an equal volume of formamide containing 5 mM EDTA and resolved on a 20% (v/v) denaturing polyacrylamide gel. The resolved nucleic acids were stained with SYBR-GOLD (Invitrogen) and gel images were obtained using a Chemidoc-MP imaging system (Bio-Rad).

### Size exclusion chromatography

The oligomerization states of A3H Hap I mutants were determined by loading 120 to 300 μg of the purified enzymes onto a Superdex 200 10/300 Increase (GE Healthcare). The running buffer contained 20 mM Tris pH 8.0, 300 mM NaCl, 10% (v/v) glycerol and 1 mM DTT. The Bio-Rad gel filtration standard set was used to generate a calibration curve from which the apparent molecular masses and oligomerization states of the enzymes were determined.

### Steady-state fluorescence polarization

The apparent dissociation constant (K_d_) values of A3H Hap II, A3H Hap VII, and A3H Hap I mutants K117E and K117E/K121E for fluorescein labeled ssDNA (118 nt) were determined using steady state fluorescence polarization to measure rotational anisotropy. Reactions (50 μL) were conducted in deamination buffer (50 mM Tris, pH 7.5, 40 mM KCl, 10 mM MgCl_2_, and 2 mM DTT) and contained 50 nM fluorescein labeled DNA or RNA and increasing amounts of A3H. A QuantaMaster QM-4 spectrofluorometer (Photon Technology International) with a dual emission channel was used to collect data and calculate anisotropy. Measurements were performed at 21°C. Samples were excited with vertically polarized light at 495 nm (6 nm band pass) and vertical and horizontal emissions were measured at 520 nm (6 nm band pass). The K_d_ was obtained by fitting to a rectangular hyperbola or sigmoidal curve equation using SigmaPlot 11.2 software.

### Cell culture and generation of stable cell lines

NCI-H1563 and MRC-5 were obtained from ATCC and cultured in RPMI-1460 supplemented with 10% FBS, 10 mM HEPES, 1mM sodium pyruvate and DMEM supplemented with 10% FBS, respectively. Lentivirus was generated by co-transfecting psPAX2, pVSV-G, and the pLVX lentiviral vector containing A3H hap I-Flag with GeneJuice (EMD Millipore) in HEK-293T cells. Media was changed after 16 hr and viral particles were harvested 48 hr post transfection. NCI-H1563 and MRC-5 were transduced with the resulting lentivirus by incubation for 16 hr in medium containing 8 ug/mL polybrene. Transduced cells were selected with 1 ug/mL puromycin for a week and maintained with 0.25 ug/mL puromycin.

### Immunofluorescence microscopy

NCI-H1563 and MRC-5 cells were treated with 2 ug/mL doxycycline for 24 h to induce A3H Hap I-Flag and fixed with 100% cold methanol for 10 min. Cells were permeabilized with 100% cold acetone for 1 min and 0.5% Triton X-100 for 10 min. Anti-Flag and anti-γH2AX antibodies (Invitrogen) in 3% BSA in 4x SSC buffer were incubated 1 hr. Primary antibodies were detected with Alexafluor-594 and Alexafluor-488-conjugated secondary antibodies (Invitrogen). Nuclei were stained with DAPI and cells were imaged using the Zeiss LSM700 system.

### qPCR

The RNA preparation, cDNA synthesis and qPCR was carried out according to Refsland et al.^91^

## Supplementary Information

**RMSD, RMSF, PCA, and correlation matrices for individual replicates**, coordinate, and parameter files for each system, deamination activity, expression and purity are provided. Source code and R scripts for energy decomposition analysis are provided as additional SI material.

## Notes

#### Summary of Updates

references cited

