## supplemental info for "Induced mutagenesis by the DNA cytosine deaminase APOBEC3H Haplotype I protects against lung cancer" (from another version of this preprint: version 1, 2020-02-28)

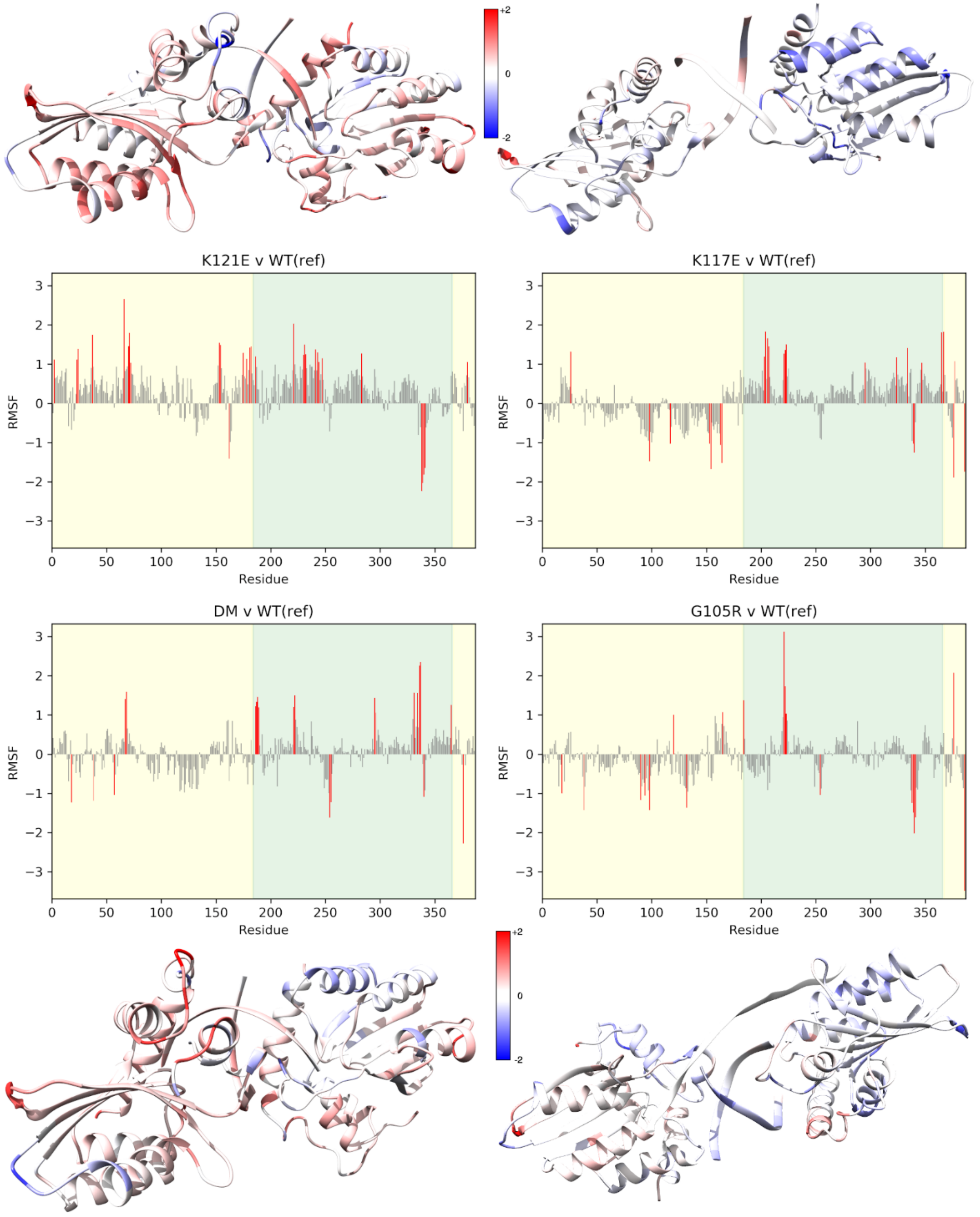


**Figure SF1**: Differences in RMSF by residue for all systems compared to the wildtype.  Residues highlighted in red change by more than 1Å.

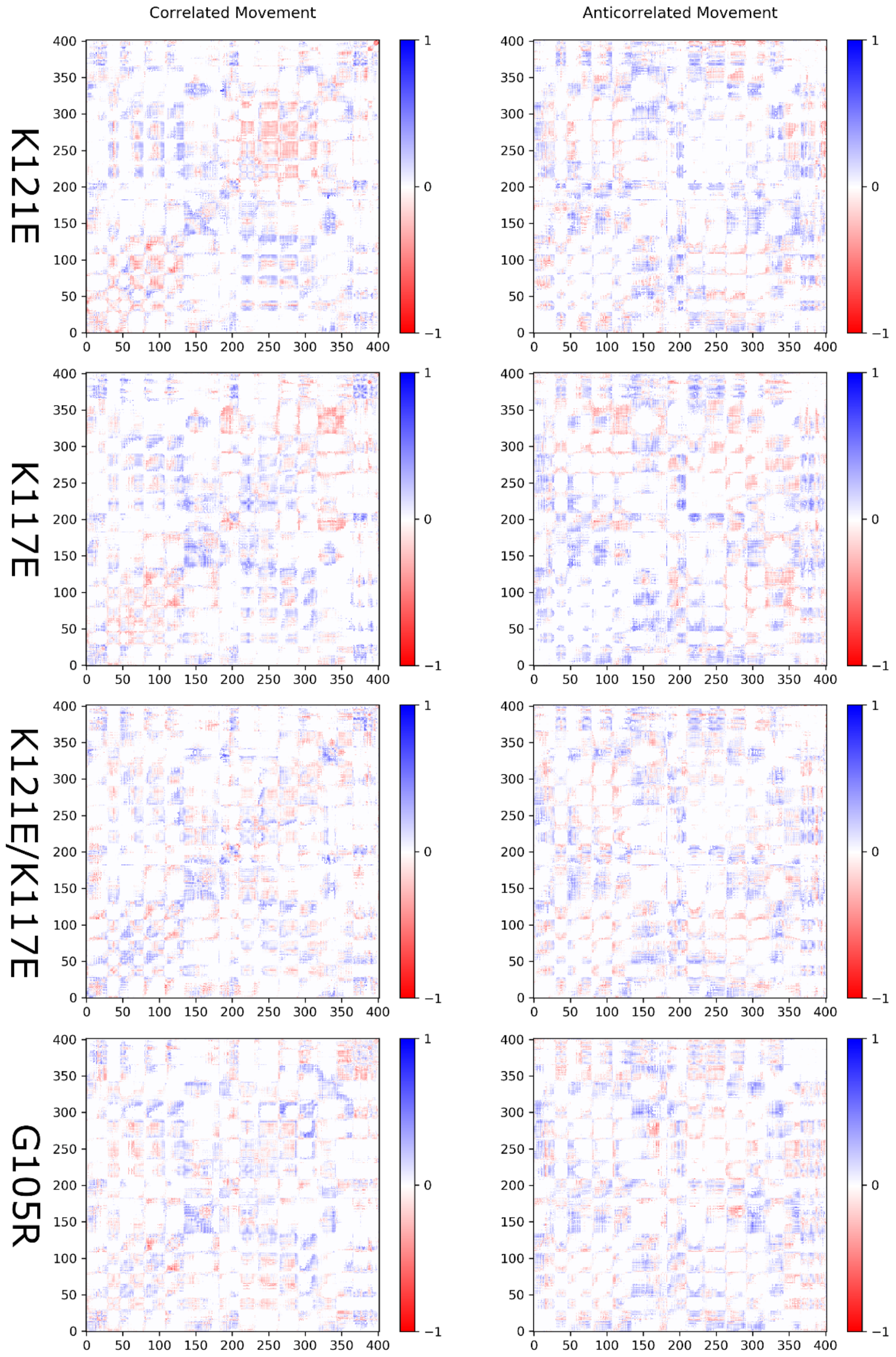


**Figure SF2**: Difference correlation matrices comparing mutant systems to wildtype. Regions of blue exhibit an increased magnitude of correlation (left) or anticorrelation (right), regions of red indicate decreased magnitude.

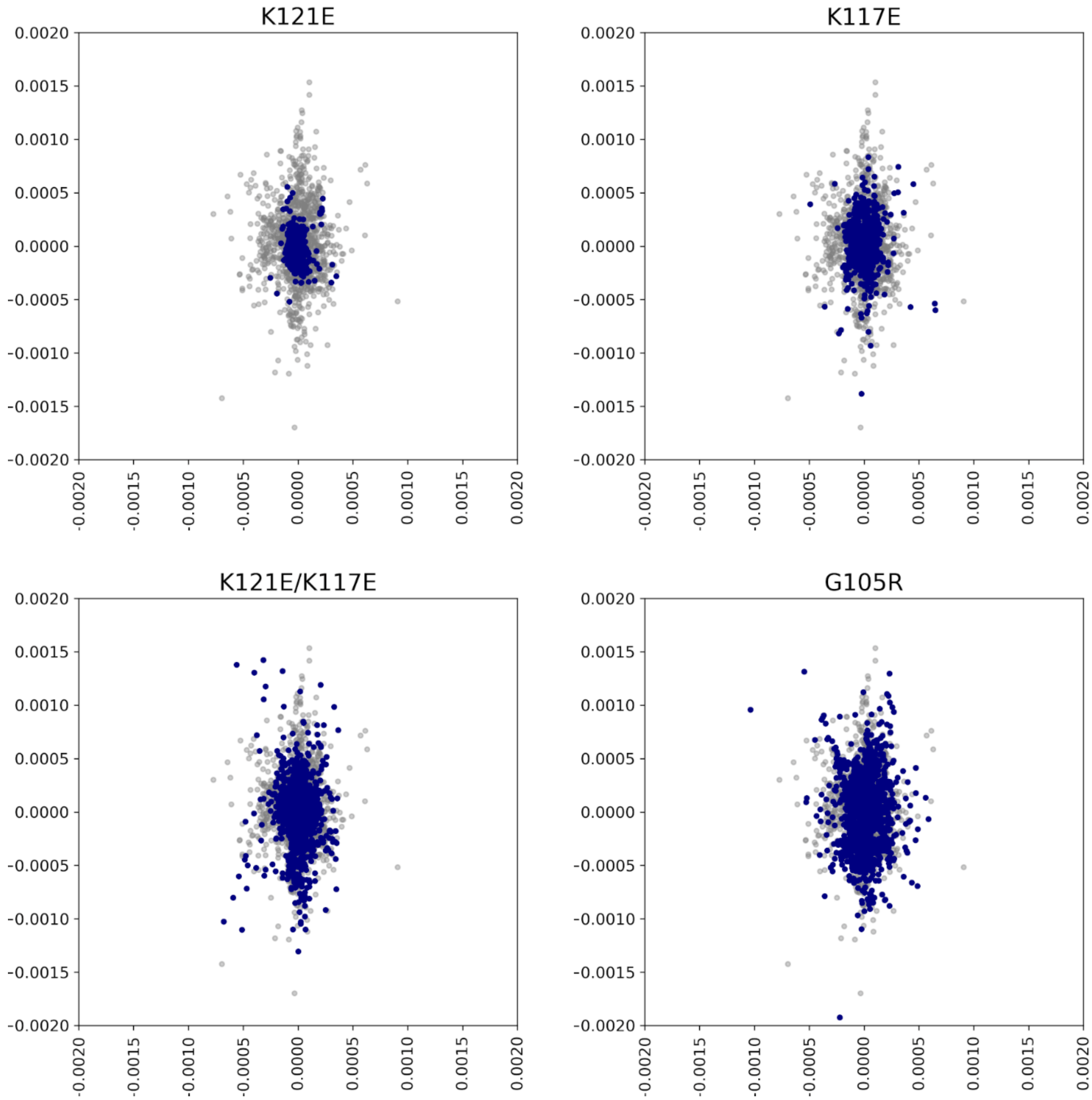


**Figure SF3**: Principle Component Analysis (PCA) of all A3H systems comparing the first two modes of each system. Points in grey are the wildtype, which is shown as a reference for the individual mutations. K121E exhibits greatly diminished scatter in the first two modes. K117E shows slightly diminished scatter. K121E/K117E rescue and G105R both exhibit similar modes of motion to the wildtype.

| | WT | K121E | K117E | K121E/K117E | G105R |
|---|---|---|---|---|---|
| K/E121 - K/E117 | 56.61% | 65.39% | 55.36% | 53.01% | 0.76% |
| K/E121 - R124 | 0.77% | 38.65% | 0.31% | 41.65% | 0.09% |
| R124 - I182 | 2.93% | 19.26% | 2.16% | 6.24% | 0.68% |
| P118-S86 | 4.46% | 47.28% | 33.70% | 26.26% | 0.00% |
| R175 - G374 | 35.27% | 4.39% | 5.19% | 44.88% | 39.77% |
| R175 - G375 | 24.78% | 6.20% | 33.75% | 35.76% | 3.23% |
| R175 - C376 | 0.00% | 10.25% | 0.00% | 0.00% | 0.68% |
| R176 - G374 | 0.06% | 30.63% | 9.15% | 0.01% | 4.86% |
| R176 - G375 | 27.23% | 44.87% | 76.64% | 76.47% | 22.92% |
| R176 - C376 | 10.10% | 10.06% | 4.85% | 87.29% | 2.77% |
| R179 - G374 | 24.88% | 0.03% | 0.00% | 0.00% | 7.21% |
| R179 - G375 | 27.72% | 23.36% | 10.06% | 60.68% | 30.01% |
| R179 - C376 | 1.71% | 52.30% | 43.21% | 84.41% | 9.69% |
| Q120 - C376 | 41.37% | 4.65% | 14.17% | 86.92% | 0.24% |

**Table ST1**: Largest contributors to hydrogen bonding interactions presented as percentage of total simulation time. These interactions form a network across a portion of the protein surface including active site residues and RNA-binding residues.

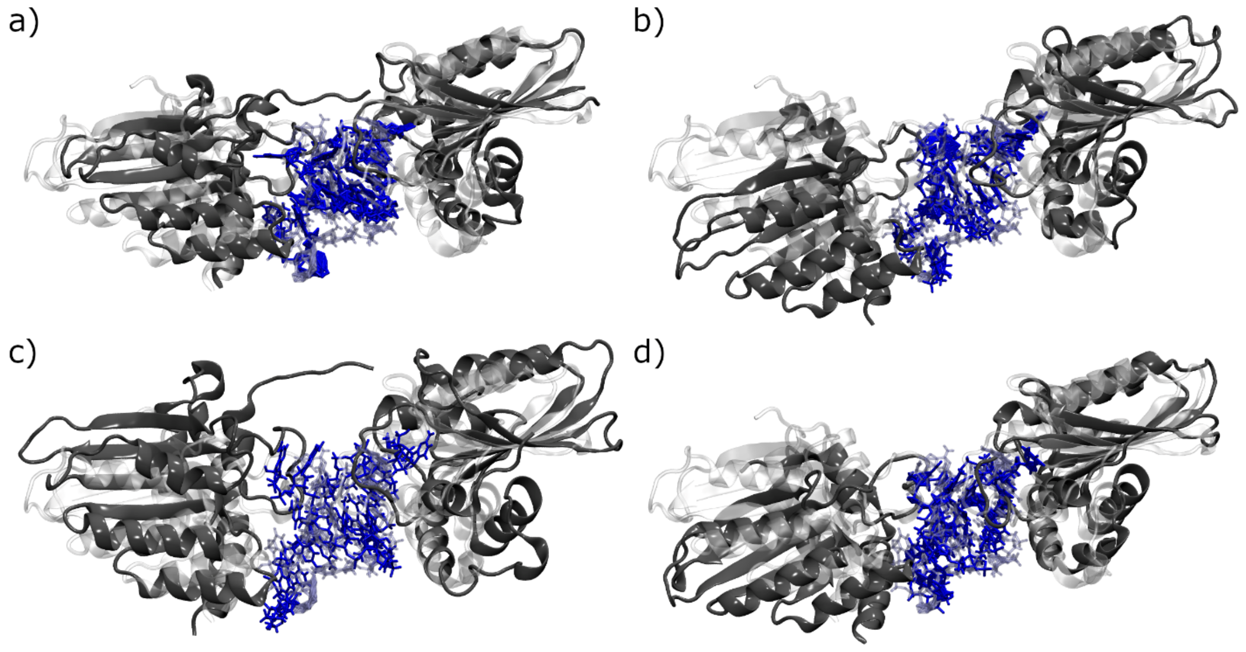


**Figure SF4**: Average structure overlay with wildtype shown as transparency for a) K121E, b) K117E, c) K121E/K117E, and d) G105R.

| | $K^+$ ions added to neutralize |
|---|---|
| WT | 12 |
| K121E | 16 |
| K117E | 16 |
| K121E/K117E | 20 |
| G105R | 10 |

**Table ST2**: Potassium Counterions added to each A3H dimer system to bring total system charge to 0.

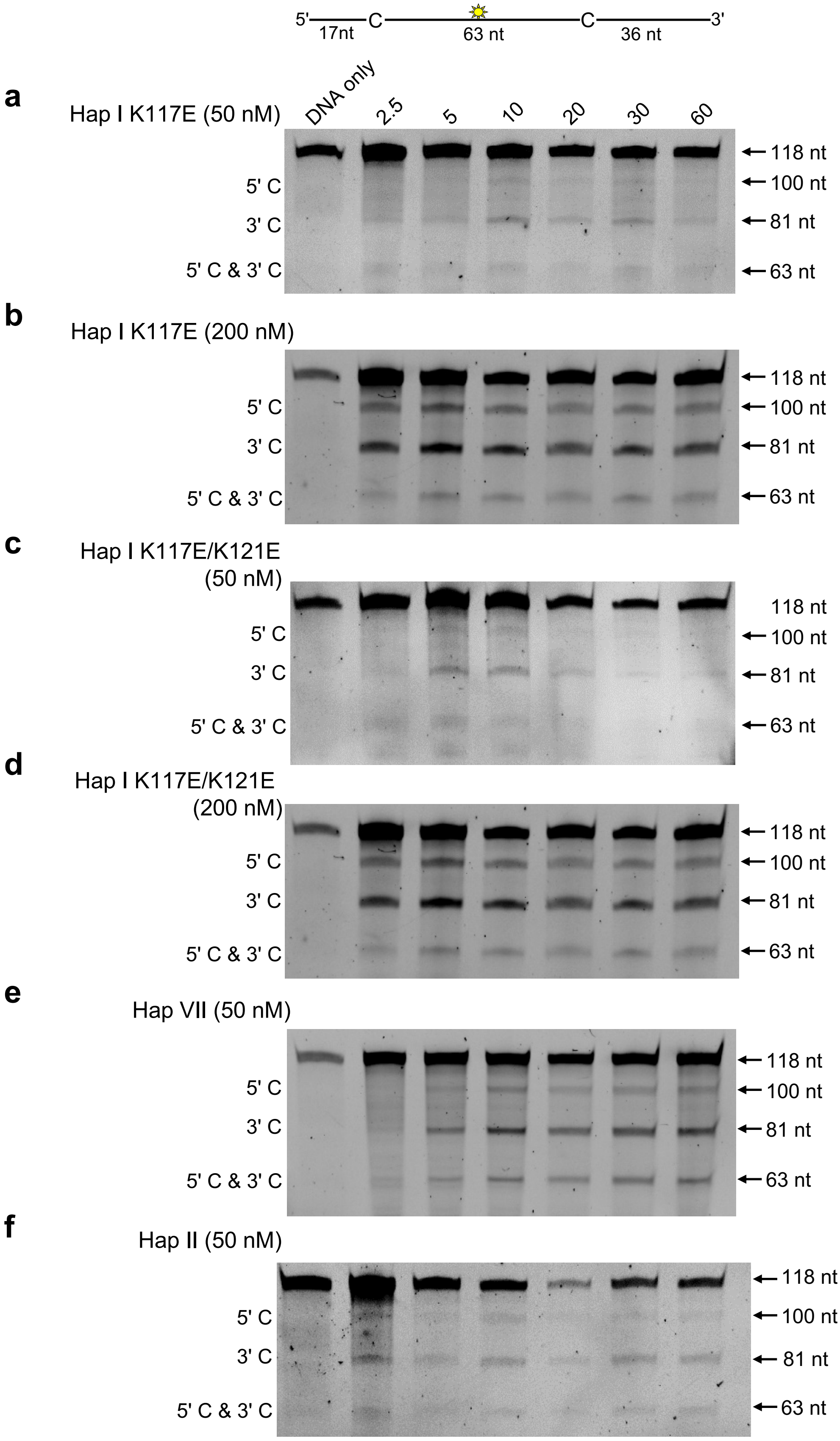


**Figure SF5: Deamination activity of A3H Hap I WT and mutants.** Time course of deamination of A3H Hap II, A3H Hap VII, and A3H Hap I WT and mutants at 50 nM or 200 nM of enzyme. Deamination was tested on a 118 nt ssDNA (100 nM) and gels analyzed for each plot were quantified and plotted as shown in Figure 2b.

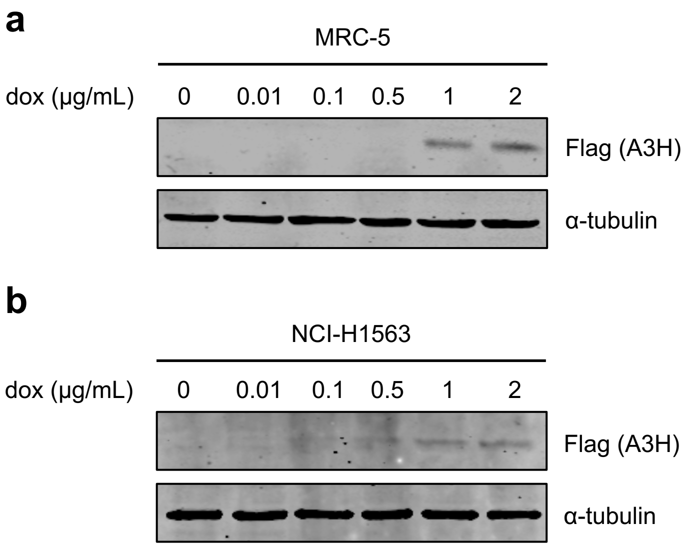


**Figure SF6: Expression of A3H Hap I-Flag in doxycycline (dox) inducible cell lines.** Titration of dox in **a)** MRC-5 and **b)** MCI-H1563 cell lines shows that expression occurs at 1 µg/mL Dox or higher.

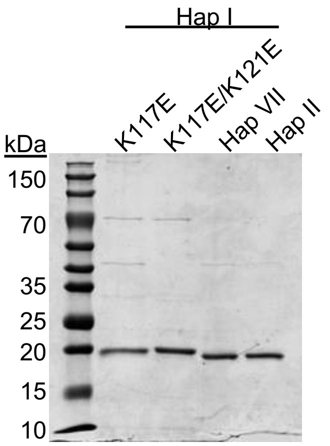


**Figure SF7: Purity of A3H enzymes.** Purified A3H enzymes were resolved by SDS-PAGE.
